# Liver FoxO1 overexpression is positively associated with the degree of liver injury in cirrhotic patients

**DOI:** 10.1101/2020.11.05.369504

**Authors:** Esther Fernandez-Galán, Silvia Sandalinas, Blai Morales-Romero, Laura Macias, Montse Pauta, Jordi Ribera, Gregori Casals, Loreto Boix, Jordi Bruix, Wladimiro Jimenez, Manuel Morales-Ruiz

**Affiliations:** Department of Biochemistry and Molecular Genetics, Biomedical Diagnostic Centre (CDB), Hospital Clinic of Barcelona; Institut d’Investigacions Biomèdiques August Pi i Sunyer (IDIBAPS), Centro de Investigación Biomédica en Red de Enfermedades Hepáticas y Digestivas (CIBERehd), Barcelona, Spain; Grupo de Trabajo de Valoración Bioquímica de la Enfermedad Hepática SEQC^ML^; Barcelona Clinic Liver Cancer Group, Liver Unit, Hospital Clínic of Barcelona; Department of Biomedicine-Biochemistry Unit, Faculty of Medicine and Health Sciences. University of Barcelona, Barcelona

**Author notes:** **Corresponding author:** Manuel Morales-Ruiz, Department of Biochemistry and Molecular Genetics, Hospital Clinic of Barcelona, 170 Villarroel Street, 08036 Barcelona, Spain.

**Keywords:** Akt, FoxO1, cirrhosis, liver regeneration

## Abstract

**Introduction:** Chronic liver disease is associated with high mortality. Liver transplantation is the definitive treatment for patients with end-stage liver disease, improving their survival and quality of life. However, chronic rejection of the graft and the imbalance between the demand and the availability of organs limit its applicability. Therefore, finding therapeutic and/or diagnostic alternatives for these patients is a priority. In this context, preclinical studies in rodents have demonstrated that Akt plays a key role in liver dysfunction. Even with all this evidence, the activation status of Akt and its downstream targets in the liver of patients with chronic hepatopathy is still unknown. Hence, the present study aims to determine the activation status of the molecules involved in the Akt signaling pathway in livers of cirrhotic patients.

**Materials and Methods:** In this study, 36 liver tissue samples from a cohort of 27 cirrhotic patients and 9 patients without cirrhosis were included. A total of 10 proteins involved in Akt/mTOR pathway (GSK3β, IGF1R, IRS1, mTOR, p70S6K, IR, PTEN, GSK3α, TSC2, and RPS6) were analyzed using a multiplex immunoassay based on Luminex® technology.

**Results:** Significant differences were found in several Akt/mTOR target proteins between the groups of cirrhotic patients vs. non-cirrhotic: FoxO1 (9.5 vs. 4.4; *p*<0.01), p-Akt (2.1 vs. 1.0; *p*<0.01), PTEN (3.061 vs. 1.877; *p*<0.05) and p70S6K (196.3 vs. 270.5; *p*<0.001). FoxO1 showed the best correlation with biochemical markers of liver injury aspartate aminotransferase and serum alanine aminotransferase (ASAT: r=0.51, *p*<0.05; ALAT: r=0.49, *p*<0.05). Moreover, the individual influence of FoxO1 on these parameters was confirmed by multiple regression analysis. It was the only enzyme in the Akt signaling pathway identified as a positive independent predictor of increased ASAT and ALAT levels.

**Conclusion:** FoxO1 is overexpressed in the liver of cirrhotic patients after partial hepatectomy. FoxO1 levels are also associated with the degree of liver injury, showing a positive correlation with current biomarkers used in clinical practice to detect liver injury.

## 1. INTRODUCTION

In Spain, most of the centers with more than 500 beds offer specific consultations for chronic liver disease complications such as hepatocellular carcinoma (HCC) (57%), cirrhosis (46%), and liver transplantation (46%). Moreover, this condition is the diagnostic category that has experienced the second-highest growth in the last five years, based on existing data (2007-2011)(1). Medical care related to liver disease is complex and burdens outpatient consultations, hospitalizations and high mortality rates. One of the most remarkable facts is that this illness affects the economically active population, and therefore impacts on the productivity of society. Besides, between 7,000 and 10,000 people die each year in Spain as a result of cirrhotic disease or its associated complications. Liver transplantation is the definitive treatment for patients with end-stage liver disease, improving their survival and quality of life. However, chronic rejection of the graft, impaired liver regeneration in these patients, and the imbalance between the demand and the availability of organs compromise the effectiveness of this therapeutic strategy.

As a result of these challenges, there is an urgent need to find new diagnostic and therapeutic alternatives for these patients, to block disease progression and prevent organ failure and associated comorbidities with the subsequent socioeconomic costs.

In this context, a significant number of studies have demonstrated that protein kinase b (Akt) is a crucial element for the control of liver function. For instance, liver cirrhosis has been associated with impaired Akt activity in experimental rat models, while the restoration of its function, using gene therapy, improved liver hemodynamics (2). Several mechanisms contribute to Akt dysregulation in liver dysfunction. Among them, the existence of an interaction between GRK2, an inhibitor of G-protein-coupled receptor signaling, and Akt-mediated eNOS activation has been reported in experimental models of liver fibrosis (3).

The beneficial effect of Akt activation on liver function was corroborated by another study conducted in an experimental model of orthotopic liver transplantation in pigs. In this case, the ischemia-reperfusion injury during transplantation is the leading cause of oxidative stress and cellular death, and this fact is clinically associated with loss of graft functionality and increased risk of graft rejection. Constitutive activation of the Akt pathway, through the forced expression of the myr-Akt variant in transduced liver grafts, restored the liver function and counteracted the apoptotic processes, increasing the viability of the graft (4). This result supports previous findings about the importance of Akt signaling in the maintenance of fundamental physiological processes of the liver.

Recent studies also suggested that Akt could have an essential function in the liver regeneration process. Several publications agree that during liver regeneration, Akt is phosphorylated (active) and prevents the apoptosis of hepatocytes by induction of cell survival signaling (5)(6). The deficiency of Akt or its activating molecules, PDK1 and PI3-K, results in decreased liver regeneration and increased mortality after partial hepatectomy (7). This is because Akt and some of its targets or activators are positive regulators of cell survival, as in the case of IR, IRS, PI3-K, and PTEN.

Additionally, Akt is a crucial mediator of cellular growth through direct regulation of mTOR. Ultimately, a study carried out using an experimental model of partial hepatectomy in FoxO1-deficient mice, demonstrated that specific inhibition of FoxO1, mediated by Akt, in hepatocytes is essential for liver regeneration (8).

Notwithstanding all this evidence supporting the fact that Akt activity has an impact on the physiology and pathophysiology of the liver; most of the studies have been conducted in experimental models in rodents. Thus, the activation status of Akt and its downstream targets in the liver of patients with chronic hepatopathy is still unknown.

This fact limits the transfer of acquired knowledge in pre-clinical studies to the development of new therapies or biomarkers. For this reason, the evaluation of the activation status of Akt and its molecular targets in patients with chronic liver disease is clinically relevant.

### OBJECTIVES

To assess the activation status of proteins involved in Akt/mTOR pathway in the liver of patients with cirrhosis. Additionally, we want to evaluate the association between Akt proteins and liver injury and loss of functionality, as well as to determine their diagnostic and prognostic value.

## 2. MATERIALS AND METHODS

### 2.1. Study design and subjects

In this retrospective study, we included specimens from patients with or without biopsy-confirmed cirrhosis who underwent therapeutic partial hepatectomy. A total of 36 liver tissue samples were analyzed, of which 27 were cirrhotic tissue samples and 9 samples of healthy tissue (non-cirrhotic group). Cirrhotic liver samples were obtained by liver resection from patients with cirrhosis associated with hepatitis C virus infection (HCV). The samples used as “non-cirrhotic group” were fragments of healthy liver tissue obtained from resections of colorectal metastasis before vascular clamping. Samples with the histological presence of tumor tissue were excluded for further analysis. The demographic characteristics of the patients included in the study are shown in **Table 1**.

**Table 1.**
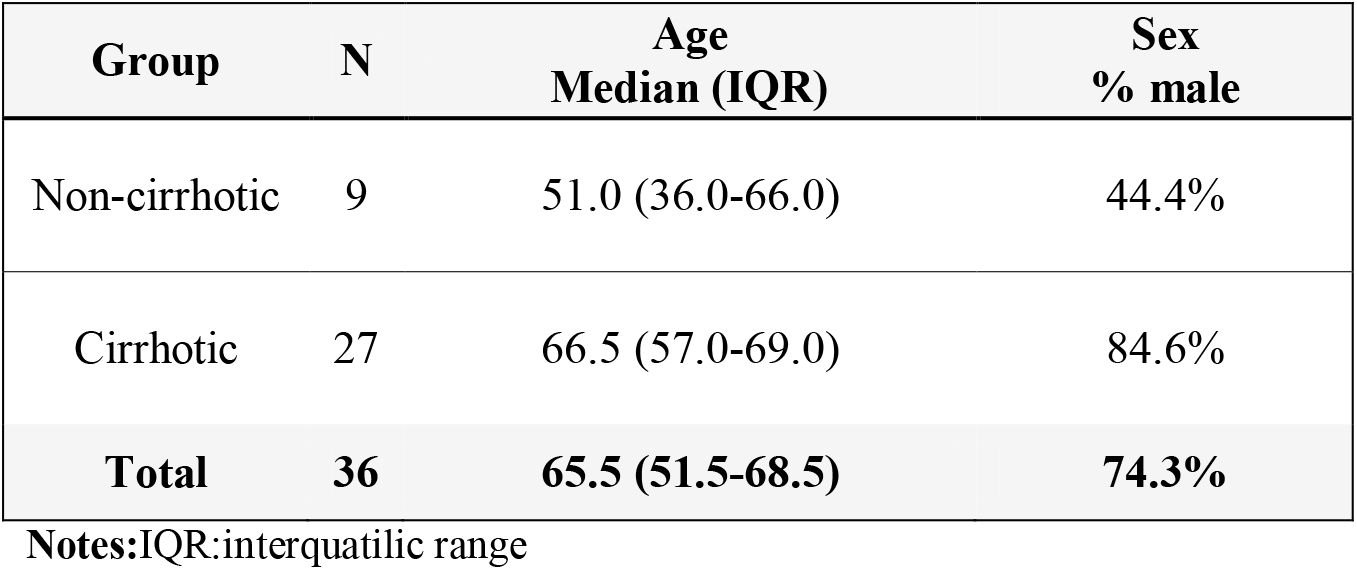
Demographic characteristics of the patients included in this study.

The present study was approved by the Hospital Clinic Ethics Committee, and informed consent was obtained from all patients. Likewise, this study was conducted in accordance with the ethical principles of the 1975 Declaration of Helsinki for medical research involving human subjects.

### 2.2. Analytical parameters of liver injury and function

In the group of cirrhotic patients, several biomarkers of liver injury and functionality were evaluated, to study their possible correlation with target proteins involved in the Akt pathway. For this purpose, biochemical markers were determined following the established clinical protocol: aspartate aminotransferase (ASAT), alanine aminotransferase (ALAT), gamma-glutamyl transferase (GGT); albumin and bilirubin. The tumor marker, alpha-fetoprotein (AFP), was also determined, among other parameters included in the hepatic panel: glucose, cholesterol, and triglycerides. All these parameters were determined in serum prior to the liver resection surgery. All the assays were carried out at the CORE laboratory of the Hospital Clínic of Barcelona, using the autoanalyzers ADVIA 2400 Chemistry System and ADVIA Centaur XP (Siemens Healthineers, Tarrytown, NY).

### 2.3. Luminex^®^ Immunoassay: analysis of proteins involved in the Akt/mTOR pathway

We determined the levels of 10 phosphoproteins involved in the Akt / mTOR signaling pathway in a total of 36 liver tissue samples. The following analytes: GSK3β, IGF1R, IRS1, mTOR, p70S6K, IR, PTEN, GSK3α, TSC2, and RPS6 were quantified using a multiplex immunoassay based on Luminex^®^xMAP^®^technology, the Akt/mTOR phosphoprotein magnetic bead panel 11-plex 96-well plate assay (Merck Millipore, Darmstadt, Germany). This assay uses paramagnetic beads (6.5 μm diameter) to fix analytes and reaction substrates, allowing the quantification of multiple analytes in a single test. Assays were performed according to manufacturer’s instructions, standards and samples were run in duplicate, the incubation step was performed overnight with shaking at 4°C (18 h, 750 rpm), and a hand-held magnetic block was used for the washing steps. Data was acquired on a Luminex MagPix 200 system (Luminex, Molecular Diagnostics, Toronto, Canada) and analyzed with the XPonent software (Luminex, Molecular Diagnostics, Toronto, Canada).

For the liver tissue analysis, it was necessary to perform a sample pre-treatment, which included the homogenization of 100 mg of tissue in a Polytron Homogenizer (Thomas Scientific). Subsequently, the total protein concentration was quantified and normalized in all samples using the Pierce BCA protein assay kit (ThermoFisher Scientific, Waltham, MA, USA). Results were expressed as median fluorescence intensity (MFI).

### 2.4. Western Blot: expression levels of Akt and FoxO1 proteins

The tissue abundance of Akt and FoxO1 proteins, as well as their degree of phosphorylation, was determined by Western Blot. Tissue lysates were prepared in a lysis buffer (Tris-HCl 20 mM [pH 7,4]) containing 1% Triton X-100, 0.1% sodium dodecyl sulfate (SDS), 50mM NaCl, 2.5 mM ethylenediaminetetraacetic acid, 1 mM Na_4_P_2_O_7_·10H_2_O, 20 mM NaF, 1 mM Na_3_VO_4_, 2 mM Pefabloc and protease inhibitor (Mini Complete, Roche, Switzerland).

Proteins were separated on a polyacrylamide gel with 10% of sodium dodecyl sulfate(SDS) (Mini Protean III; Bio-Rad, Richmond, CA), and they were transferred for 2 hours at 48°C to nitrocellulose membranes of 0.45 μm. After blocking, the membranes were incubated at 48°C overnight with the following antibodies: rabbit anti-FoxO1 (1:1000; Cell Signaling), rabbit anti-Akt (1:5000; Cell Signaling), rabbit anti-phospho-Akt (Ser473) (1:5000; Cell Signaling) and anti-tubulin antibody (1:5000 Cell Signaling). Next, the membranes were incubated with peroxidase-conjugated secondary antibodies at 1: 5000 dilution (GE Healthcare) for 1 hour at room temperature. The respective bands were visualized using Luminata Forte Western HRP Substrate (Millipore) e Image-Quant LAS 4000 (GE Healthcare). Densitometry analysis of the gels was performed using the Image J software (version 1.37). Results were expressed in densitometric relative units (DRU), as a result of the densitometric ratios of p-Akt/Akt and FoxO1/tubulin.

### 2.5. Statistical analysis

Continuous variables were expressed as medians ± interquartile range (IQR). Differences in quantitative variables among groups were evaluated using the Mann-Whitney U test. The degree of correlation between variables was studied using the Pearson or the Spearman coefficient, depending on the variable distribution. For linear correlation and linear regression analysis, the FoxO1 variable was logarithmically transformed (LnFoxO1) to correct the skewness of the distribution. The parameters that showed a significant or near significant correlation with analytical parameters of liver injury and function, were selected for the multiple regression analysis. The multivariate statistical analysis performed was linear regression. The validity of the resulting models was evaluated plotting residuals or Pearson residual against adjusted values and plotting the influence of each observation on the adjusted response according to Cook’s distance. All statistical analyses were performed using public libraries from “The Comprehensive R Archive Network” (CRAN; http://CRAN.R-project.org) rooted in the open-source statistical computing environment R, version 3.1 (http://www.R-project.org/). The software “GraphPad Prism version 8.0.2” (GraphPad Software, La Jolla California USA) was used for graphic representations. *p*-value ≤ 0.05 was considered significant.

## 3. RESULTS

### 3.1. Analytical parameters of liver injury and function

All cirrhotic patients showed aminotransferases (ALAT and ASAT) and GGT levels compatible with liver cirrhosis, as shown in **Table 2**. These increases in biochemical markers of liver injury were only observed in the serum samples from cirrhotic patients and not in the samples from the non-cirrhotic patients (data not shown).

**Table 2.** Analytical parameters determined in the serum of cirrhotic patients.

| Parameters | N | Median (IQR) | Reference range |
| --- | --- | --- | --- |
| **Markers of liver injury and function** |  |  |  |
| ASAT | 25 | **71.0 (52.0-114.0)** | 5-40 IU/L |
| ALAT | 25 | **113.0 (72.0-165.0)** | 5-40 IU/L |
| GGT | 25 | **71.0 (38.0-140.0)** | 5-40 IU/L |
| Albumin | 24 | 43.0 (39.5-45.0) | 34-48 g/L |
| Bilirubin | 22 | 0.8(0.6-1.0) | < 1.2 mg/ dL |
| **Others** |  |  |  |
| Glucose | 25 | 99.0 (87.0-119.0) | 65-110 mg/dL |
| Cholesterol | 24 | 164.5 (149.5-180.5) | ≤ 200 mg/dL |
| Triglycerides | 24 | 99.5 (86.5-135.0) | < 150 mg/dL |
| Alfa-fetoprotein | 20 | **12.5 (7.5-49.0)** | < 10 ng/mL |

**Notes:** Variables are expressed as median and interquartile range (IQR). Values outside the range are in bold. IU: International Units, ALAT: alanine aminotransferase, ASAT: aspartate aminotransferase, GGT: gamma-glutamyl transferase. N: number of patients in which it was possible to determine the parameter.

### 3.2. Comparative study: Intrahepatic expression of Akt pathway proteins in cirrhotic patients vs. non-cirrhotic

Determination of protein expression levels (FoxO1 and Akt) in liver tissues, were carried out by Western blot analysis. The results obtained showed increased levels of p-Akt (active form) in the group of cirrhotic patients compared to the non-cirrhotic group. These differences were statistically significant (*p*<0.01), see **Figure 1A**. By contrast, no statistically significant differences were found in the Akt levels between both studied groups. Levels of FoxO1, which is negatively regulated by Akt, were represented in **Figure 1B**. The median of FoxO1 levels was significantly higher in the cirrhotic group compared to the non-cirrhotic group (*p*<0.01).

**Figure 1.**
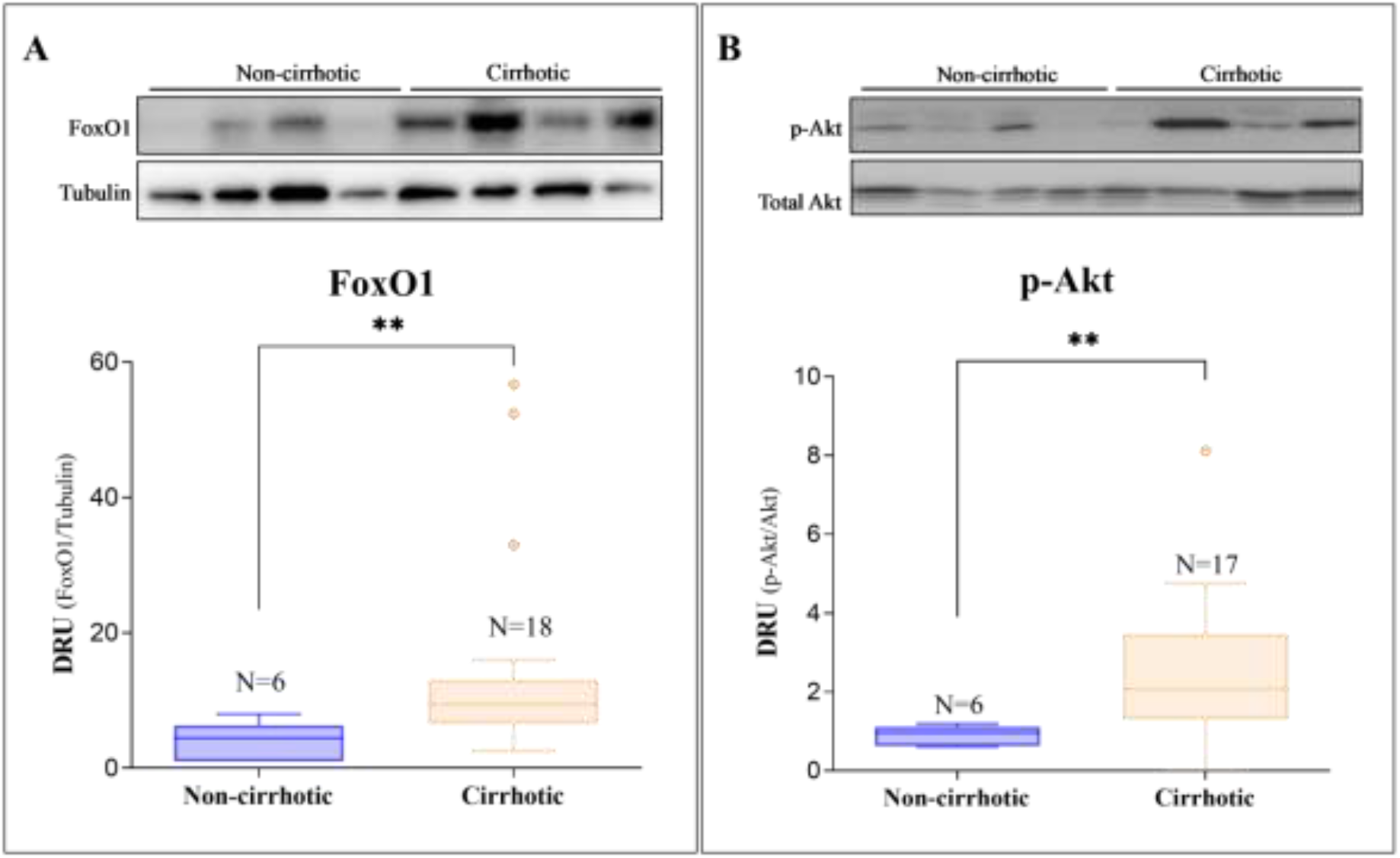
Expression levels of FoxO1 and p-Akt proteins in liver of cirrhotic patients vs. non-cirrhotic. Box plot comparing FoxO1 (A) and p-Akt (B) levels expressed as densitometric relative units (DRU). DRU were calculated as the ratio between the specific bands FoxO1/tubulin and p-Akt/Akt quantified with the software ImageJ A representative image of Western blot analyses is shown at the top of each figure. Statistically significant differences (**p <0.01) were observed when comparing the median of both proteins between groups: **FoxO1**: median = 4.44 (non-cirrhotic) vs. 9.50 (cirrhotic), **p-Akt**: median = 0.97 (non-cirrhotic) .vs. 2.07 (cirrhotic).

The other target proteins from the Akt pathway were quantified using a multiple immunoassay panel based on the technological platform Luminex^®^. Among all the proteins analyzed --GSK3β, IGF1R, IRS1, mTOR, p70S6K, IR, PTEN, GSK3α, TSC2 y RPS6--statistically significant differences were observed in 2 proteins: p70S6K and PTEN. The PTEN median levels were increased in the cirrhotic group compared with the non-cirrhotic (*p*<0.05). On the contrary, p70S6K levels were decreased in cirrhotic patients compared to the group of patients without cirrhosis (*p* <0.01), as represented in **Figure 2**. For the rest of the variables studied, no significant differences were found when comparing the median between both groups (**Table 3**).

**Table 3.**
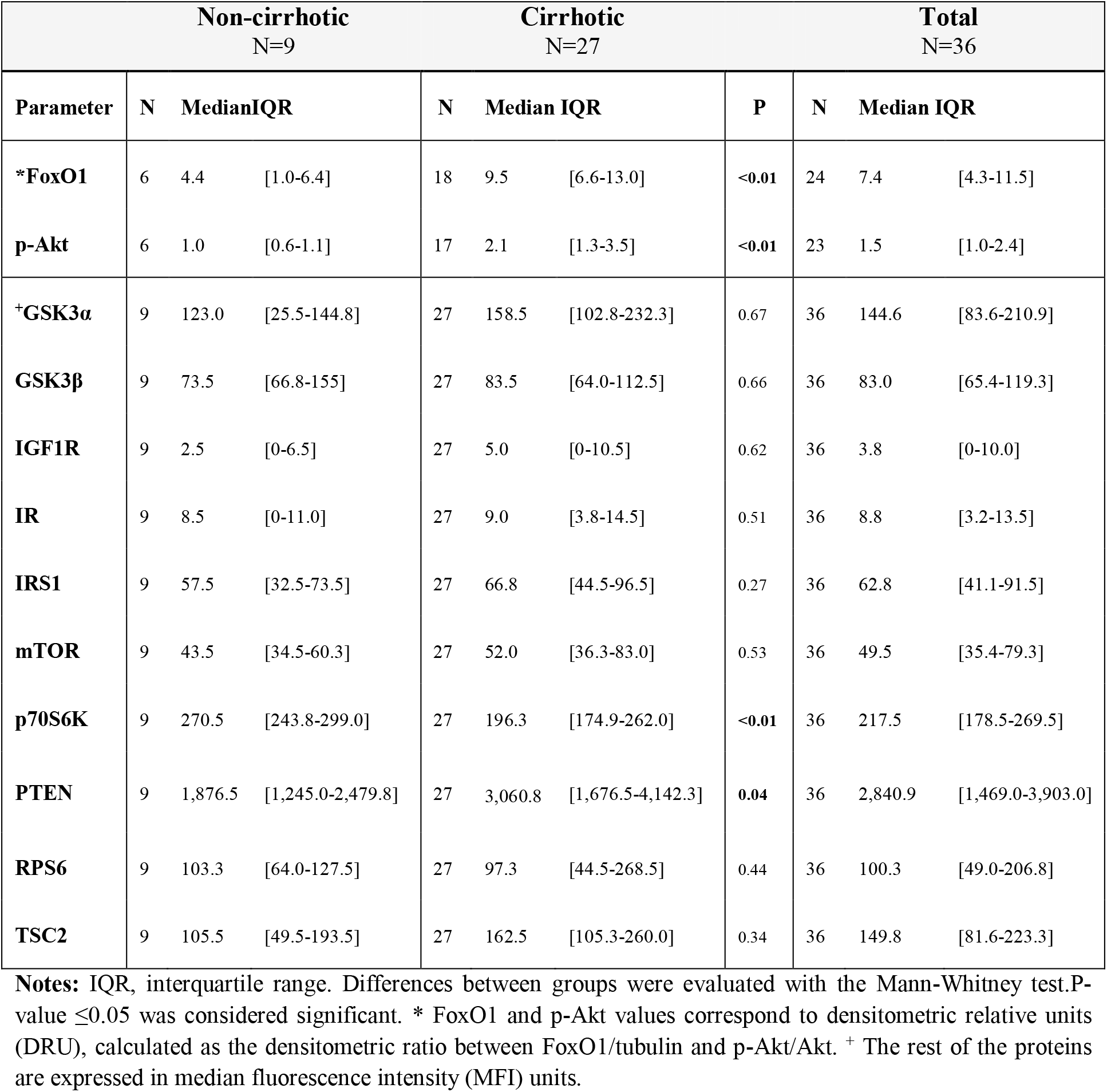
Comparative study of 12 target proteins from the Akt pathway.

**Figure 2.**
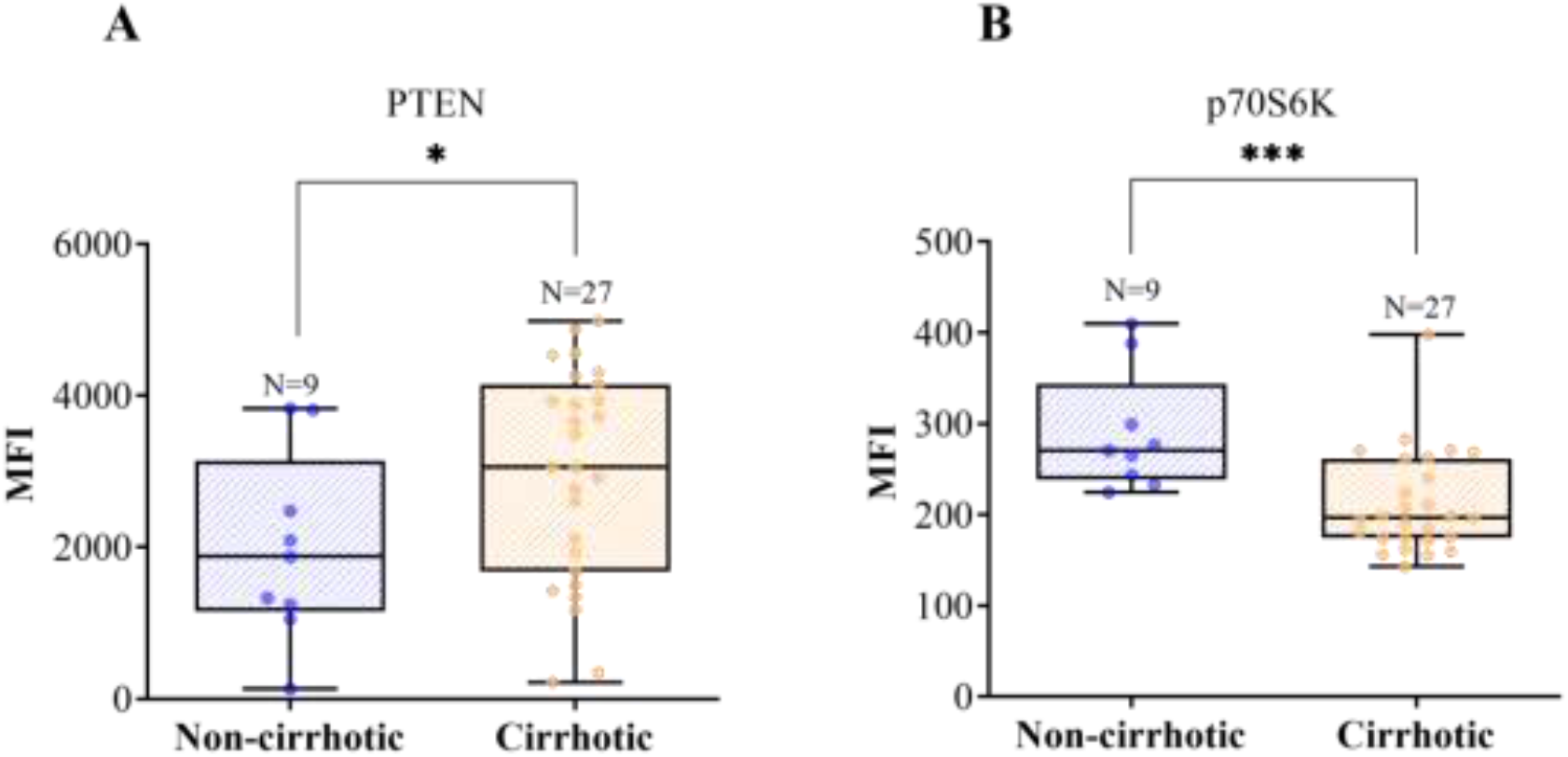
Box plot representative of p70S6K (A) and PTEN (B) levels in both groups. The levels of each protein in liver tissue are expressed in MFI (median fluorescence intensity). Box-plot and also the individual values for each patient are represented. **A.** PTEN levels are higher in cirrhotic patients (median=3061, N=27) compared with patients without cirrhosis (median=1877, N=9). These differences are statistically significant (* *p* <0.05). **B.** The group of cirrhotic patients presents lower levels of p70S6K (196.3; N=27) compared with the group of non-cirrhotic patients (270.5; N=9). These differences are statistically significant (*** *p* < 0.001).

### 3.3. Correlation between Akt pathway proteins and analytical parameters of liver injury and function

The association between all the studied proteins from the Akt pathway and the analytical parameters of functionality and liver injury has been studied. **Figure 3** shows the correlation matrix resulting from this analysis. The degrees of linear correlation, as well as its statistical significance, were calculated according to the Pearson correlation coefficient. Variables that did not fit a normal distribution were excluded from this matrix, and their relationship was evaluated individually using the Spearman correlation coefficient. The results obtained revealed that FoxO1 presented the best correlation with liver injury tests. The FoxO1 protein expression levels and serum concentration of both transaminases showed a positive (moderate-high) and statistically significant correlation: ASAT (r = 0.51, *p*=0.036) and ALAT (r = 0.49, *p* =0.049). The linear correlation and the corresponding fitted regression line between FoxO1 and the transaminases are detailed in **Figure 3B**.

**Figure 3.**
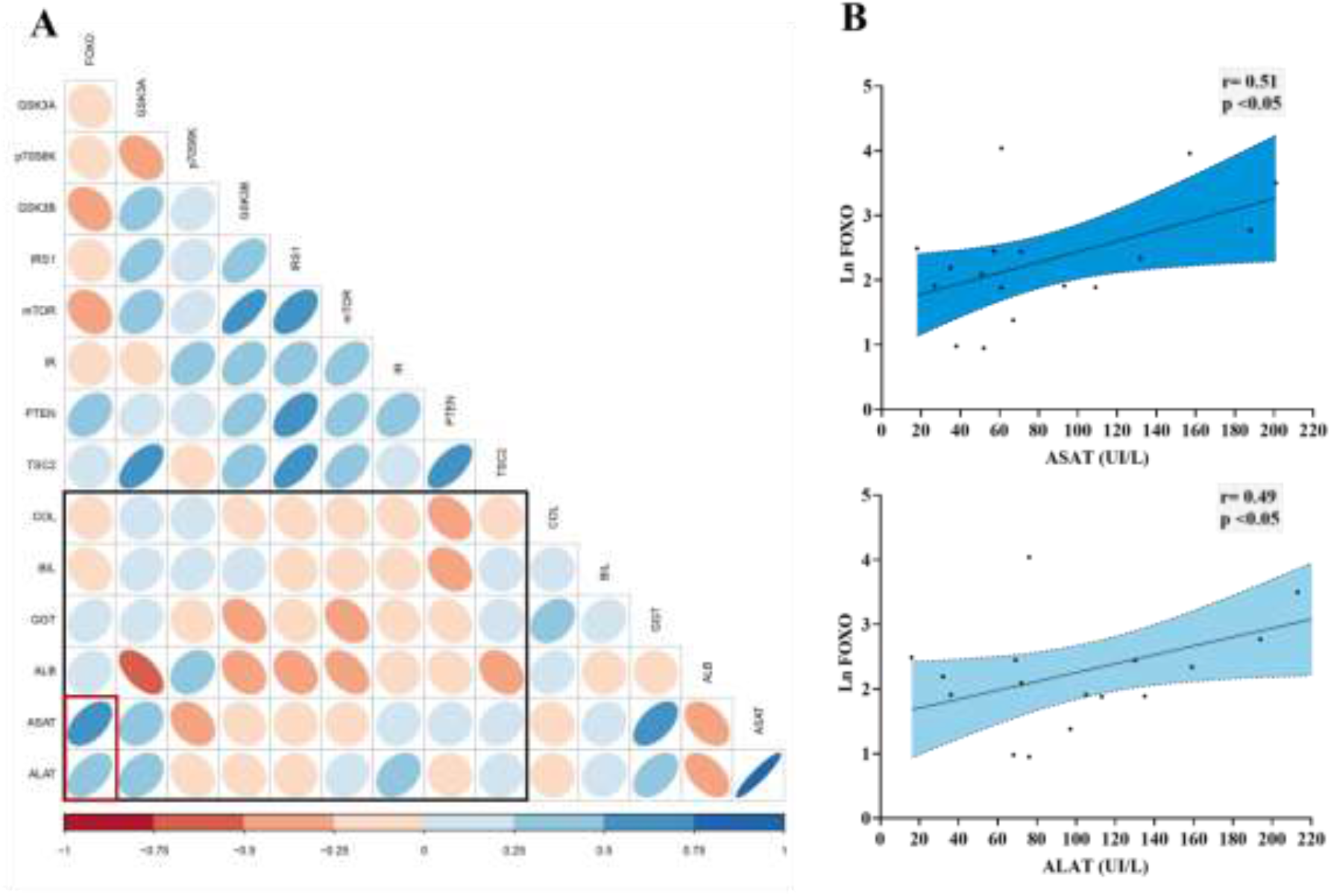
Correlation matrix between target proteins involved in the Akt pathway and analytical parameters of liver injury and function. **A:** All possible correlations between the variables studied are shown. The black rectangle highlights the correlations of interest: The most relevant correlations between FoxO1 and transaminases are inside the red box. The color scale indicates whether it is a positive (blue) or negative (red) correlation. Color intensity and the size of the ellipses are proportional to the correlation coefficients (Pearson). **B:** The scatter dot plot indicates that there is a positive correlation between levels of FoxO1 protein in liver tissue and the concentration of transaminases (ASAT and ALAT) in serum. These correlations are statistically significant, with Pearson’s correlation coefficients: r=0.51 for ASAT and r=0.49 for ALAT. The corresponding regression line is also represented; the shaded area indicates the 95% confidence intervals.

### 3.4. Regression analysis to explain the influence of FoxO1 on biochemical markers of liver injury

FoxO1 and GSK3α proteins showed a statistically significant correlation with the concentration of both transaminases in serum: FoxO1 with ASAT (r=0.51; *p* = 0.036) and ALAT (r=0.49; *p*=0.049) and GSKA with ASAT (r=0.42; *p*=0.039). For this reason, both Akt targets were included in the multivariate analysis, which showed that both parameters were positive independent predictors of ASAT, explaining 37% of the variance (R^2^-adj=0.37; F=5.75; *p*=0.015). In the case of multivariate linear regression analysis for the dependent variable ALAT, only FoxO1 was identified as a positive independent predictor explaining 18% of the variance (R^2^-adj=0.18; F=4.61; *p*=0.049) (**Table 1 Supplementary**).

## 4. DISCUSSION

The PI3K/AKT signaling pathway is crucial for energy homeostasis, cell growth, and survival. To date, studies in rodents have extensively shown that Akt plays a key role in liver regeneration as well as in normal liver function (9). The actions of the Akt pathway on the cell cycle are closely linked to FoxO1 inhibition, a member of the Foxo (forkheadbox) transcription factors. These factors are necessary for multiple functions in the liver: they regulate the stress response, adaptation to fasting, and cell proliferation (10). It is well known that Akt inactivates FoxO1, by phosphorylation, which leads to its nuclear exclusion and thus inhibits its pro-apoptotic signals (11). For this reason, the Akt-FoxO1 interaction has been identified as the most important mechanism in liver regeneration. In summary, the maintenance of Akt activity and consequent inactivation of FoxO1 is associated with better functionality and regenerative capacity of the liver. Therefore Akt and FoxO1 have been proposed as promising therapeutic targets for cirrhosis and other chronic liver diseases. However, there is currently insufficient evidence on the degree of activation of these molecules in cirrhotic patients, which hinders their possible translation to the clinic.

Our study is the first to comprehensively assess the activation status of Akt signaling pathway molecules in livers of cirrhotic patients. Previously, we demonstrated in experimental models of cirrhotic rats that the enzymatic activity of Akt is decreased in the dysfunctional liver (2). However, in the present study carried out in cirrhotic patients after partial hepatectomy, we have shown that Akt and its target FoxO1 are overexpressed in liver cirrhosis. These conflicting results should be interpreted cautiously since these discrepancies could be attributed to other factors such as comorbidities present in cirrhotic patients or the etiology of the disease. The latter is not present in the animal model, in which cirrhosis is induced by carbon tetrachloride inhalation. In our cohort of patients with cirrhosis, the baseline etiology in all cases was HCV infection. The relationship between the Akt pathway and viral replication processes in chronic HCV infection is still unknown. In vitro studies suggest that the NS5A viral protein can activate the PI3K-Akt pathway, thus inhibiting cellular apoptosis (12). Some authors have described this relationship as an evolutionary mechanism used by most DNA mammalian viruses to promote cell survival and ensure their replication (13). The results of this study highlight the importance of validating the pre-clinical results obtained for the molecules involved in the Akt pathway in humans. The most relevant finding of the present study is that FoxO1 is overexpressed in the liver of cirrhotic patients. This finding is in line with previous research showing FoxO1 overexpression in liver tissue from patients with another chronic liver disease of different etiology, non-alcoholic steatohepatitis (14).

FoxO1 overexpression is clinically relevant, considering that most of the patients diagnosed with HCC present underlying cirrhosis (15). Currently, surgical resections of these tumors or metastatic lesions in cirrhotic patients remain a significant clinical challenge due to the associated morbidity and mortality (16). In cirrhotic patients, the regeneration capacity of hepatocytes after surgery is reduced, limiting the applicability of therapeutic resection in situations such as the treatment of HCC (17). This observation is consistent with the hypothesis that the higher the expression of FoxO1, the lower the liver’s regenerative capacity, as previously demonstrated in a murine model of liver regeneration caused by partial hepatectomy (8). Furthermore, it is coherent with the fact that in our study, FoxO1 levels were significantly correlated with the degree of liver damage. Our statistical model revealed that FoxO1 is the only, of all molecules studied, that has proven to be an independent predictor of ALAT and ASAT levels in serum. Therefore, considering both these results and our previous preclinical studies, FoxO1 overexpression could be one of the underlying mechanisms that limit liver regeneration in cirrhotic tissue. The demonstration of this hypothesis would be of great importance since the intrahepatic levels of FoxO1 could be an indicative factor of individual liver regenerative capacity and, therefore, a marker with prognostic value for patients with cirrhosis and HCC. In this context, understanding the mechanisms of induction and regulation of liver regeneration is needed to open the way for the identification of new therapeutic targets. This would favor the development of therapies based on the restoration of liver functionality, improving the prognosis of patients with advanced liver disease.

Our results support the need to continue this line of research to achieve a better understanding of the role of Akt-FoxO1 in cirrhosis. Carrying out prospective observational studies would be an appropriate strategy to determine the status of this pathway in the cirrhotic population with different etiologies. These future studies would provide evidence to support the view that selective modulation of Akt or its targets is a viable strategy with beneficial effects for the management of patients with liver disease.

## Abbreviations

Akt: Protein kinase B
HCC: Hepatocellular carcinoma
eNOS: Endothelial nitric oxide synthase
FoxO1: Forkhead box protein O1
GSK3α: Glycogen synthase kinase 3 alpha
GSK3β: Glycogen synthase kinase 3 beta
GRK2: G protein-coupled receptor kinase 2
IR: Insulin receptor
IRS1: Insulin receptor substrate 1
MFI: Median Fluorescence Intensity
mTOR: Mammalian target of rapamycin.
PDK1: Phosphoinositide-dependent protein kinase 1
PI3-K: Phosphoinositide 3-kinase
PTEN: Phosphatidylinositol 3,4,5-trisphosphate 3-phosphatase
RPS6: Ribosomal protein S6
TNFα: Tumor necrosis factor alpha
TSC2: Tuberous sclerosis complex 2
p70S6K: Ribosomal protein S6 kinase beta-1
DRU: Densitometric relative units

## Acknowledgments

This study was funded by the Ministerio de Economía y Competitividad (SAF2016-75358-R to MM-R, co-funded by FEDER). Ciberehd is funded by Instituto de Salud Carlos III.

