## Supplemental table 1 for "Liver FoxO1 overexpression is positively associated with the degree of liver injury in cirrhotic patients"

**SUPPLEMENTARY MATERIAL**

**Tabla 1.** Multiple regression analysis results.

| Dependent variable **ASAT** | $F(2.14)=5.75$ $R^2=0.45$ $R^2$ -adj=0.37 $p=0.015$ | | | | |
|---|---|---|---|---|---|
| | Coeff. | Err. Std | β | *P*-value | 95% Confidence interval |
| Constant | -52.98 | 41.72 | | | -142.46 / 36.51 |
| FoxO1 | 34.07 | 12.41 | 0.54 | 0.016 | 7.47 / 60.69 |
| GSK3A | 0.38 | 0.17 | 0.44 | 0.045 | 0.01 / 0.75 |
| Dependent variable **ALAT** | $F(1.15)=4.61$ $R^2=0.24R^2$-adj=0.18 $p=0.049$ | | | | |
| | Coeff. | Err. Std | β | *P*-value | 95% Confidence interval |
| Constant | 28.20 | 39.25 | | | -55.45 / 111.86 |
| FoxO1 | 34.20 | 15.96 | 0.49 | 0.049 | 0.25 / 68.30 |
